# Improper Localization of the OmcS Cytochrome May Explain the Inability of the *xapD*-Deficient Mutant of *Geobacter sulfurreducens* to Reduce Fe(III) Oxide

**DOI:** 10.1101/114900

**Authors:** Kelly A. Flanagan, Ching Leang, Joy E. Ward, Derek R. Lovley

## Abstract

Extracellular electron transfer through a redox-active exopolysaccharide matrix has been proposed as a strategy for extracellular electron transfer to Fe(III) oxide by *Geobacter sulfurreducens,* based on the phenotype of a *xapD*-deficient strain. Central to this model was the assertion that the *xapD*-deficient strain produced pili decorated with the multi-heme *c*-type cytochrome OmcS in manner similar to the wild-type strain. Further examination of the *xapD-*deficient strain with immunogold labeling of OmcS and transmission electron microscopy revealed that OmcS was associated with the outer cell surface rather than pili. PilA, the pilus monomer, could not be detected in the *xapD*-deficient strain under conditions in which it was readily detected in the wild-type strain. Multiple lines of evidence in previous studies have suggested that long-range electron transport to Fe(III) oxides proceeds through electrically conductive pili and that OmcS associated with the pili is necessary for electron transfer from the pili to Fe(III) oxides. Therefore, an alternative explanation for the Fe(III) oxide reduction phenotype of the *xapD-deficient* strain is that the pili-OmcS route for extracellular electron transport to Fe(III) oxide has been disrupted in the *xapD*-deficient strain.

## Introduction

The mechanisms for Fe(III) oxide reduction in *Geobacter* species are of interest because *Geobacter* species are abundant in diverse environments in which Fe(III) oxide reduction is an important biogeochemical process^1^. Mechanisms for Fe(III) oxide reduction in *Geobacter* species have been studied most intensively in *Geobacter sulfurreducens* because it was the first *Geobacter* species for which tools for genetic manipulation were developed^2^.

Two models have been proposed for final steps in Fe(III) oxide reduction by *G. sulfurreducens*, both of which incorporate the previous finding that *Geobacter* species need to be in direct contact with Fe(III) oxides in order to reduce them^3^. In the exopolysaccharide matrix model^4^, redox-active moieties, such as *c*-type cytochromes, embedded in a exopolysaccharide matrix, transfer electrons to Fe(III) oxide that come into contact with the exopolysaccharide matrix. In the electrically conductive pili (e-pili) model, electrons are transported from the cell along e-pili and the pili-associated multi-heme *c*-type cytochrome OmcS facilitates electron transfer from the e-pili to the Fe(III) oxides^1,5^.

The e-pili model has been proposed as the simplest explanation for the findings that: 1) pili are required for Fe(III) oxide reduction^6^; 2) *G. sulfurreducens* pili are highly conductive^6-11^; 3) OmcS, which is not required for electron conduction along the pili^6,7,11,12^, is required for Fe(III) oxide reduction^13^; and 4) *G. sulfurreducens* strains Aro-5^8^ and PA^14^, which each express pili with low conductivity but with OmcS properly localized on the pili, are highly impaired in Fe(III) oxide reduction. Consistent with this model is the observation that charge injected into wild-type *G. sulfurreducens* pili propagates along the pili and into the pili-associated cytochrome^9^.

The exopolysaccharide matrix model^4^ was proposed after the discovery that deletion of *xapD* (gene GSU1501), which is necessary for ~50% of exopolysaccharide production, yielded a strain with diminished capacity for Fe(III) oxide reduction^15^. An important factor in interpreting the phenotype of the *xapD*-deficient strain was the suggestion that the mutant still produced pili with attached cytochromes. However, the attachment of cytochromes to the pili was only indirectly inferred^15^ based on findings that protein preparations sheared from the outer surface of wild-type and *xapD*-deficient cells both contained: 1) a large amount of a heme-staining protein with the same molecular weight as OmcS and 2) similar bands of a 6 kDa protein in which PilA was the dominant sequence “(data not shown)”. Given the importance of verifying that OmcS was properly localized on the e-pili in order to interpret the Fe(III) oxide reduction phenotype of the *xapD*-deficient strain, the localization of OmcS was evaluated in more detail. The results demonstrate that deleting genes for outer surface components of *G. sulfurreducens* can have pleiotropic effects that must be considered when developing models for extracellular electron transport.

## Materials and Methods

### Source of organism and culturing methods

The *G. sulfurreducens* strains investigated were the previously described^16^ wild-type strain PCA (ATCC 51573) and the *xapD*-deficient strain^15^. Cultures were routinely grown under strict anaerobic conditions as previously described^2^ at 25° C in NBAF medium with 15 mM acetate as the electron donor and 40 mM fumarate as the electron acceptor. Studies were conducted with cells harvested in late-log to early stationary phase because abundant OmcS and pili are produced during this phase of growth on fumarate^17^.

### Western blot analysis of OmcS and PilA

The loosely bound outer surface protein fractions of the wild-type and *xapD*-deficient strains of *G. sulfurreducens* were isolated as previously described^13^. Outer surface protein preparations (10 μg) were separated with 12% SDS-PAGE and blotted onto a PVDF membrane with a semidry transfer cell (Bio-Rad). OmcS was detected with the previously described antisera^17^ and developed with NBT/BCIP. The cellular content of PilA monomers was evaluated in triplicate samples of wild-type and *xapD*-deficient cultures harvested at early stationary phase. Whole cell lysates (10 μg) were separated with 15% SDS-PAGE and blotted onto PVDF. PilA content was detected with Western blot and chemiluminescence, as previously described^18^.

## Immunogold Labeling and Transmission Electron Microscopy

OmcS was localized with immunogold labeling and transmission electron microscopy as previously described^17,19^. Briefly, cells were harvested with centrifugation, placed on copper grids, exposed to rabbit-raised OmcS antibodies, washed, incubated with anti-rabbit IgG conjugated with 10 nm gold secondary antibody, and then examined with transmission electron microscopy. For localization of OmcS in ultrathin sections, stationary phase cells were fixed (2% paraformaldehyde and 0.5% glutaraldehyde in 50 mM PIPES pH 7.2) for an hour, then washed, dehydrated, and embedded in LR white for sectioning and then immunogold labeled as previously described^19,20^. These experiments were conducted twice using different cultures, each time with technical replicates to qualitatively observe reproducibility. Over 40 different fields of view were obtained for each strain.

## Results and Discussion

### Western blot analysis of OmcS and PilA

Western-blot analysis of OmcS in outer surface protein preparations demonstrated that the *xapD*-deficient strain produced quantities of OmcS comparable to the wild-type strain (Fig 1A). An additional band of slightly higher molecular weight that reacted with the OmcS antibody was recovered from the *xapD*-deficient mutant, but not in the wild-type strain (Figure 1A). This result raises the possibility that some of the OmcS protein may be modified in the *xapD*-deficient mutant in a manner not observed in the wild-type strain. Regardless of the presence of this second band, these results confirm the previously reported^15^ production of OmcS in the *xapD*-deficient mutant and its localization at the outer cell surface.

**Figure 1.**
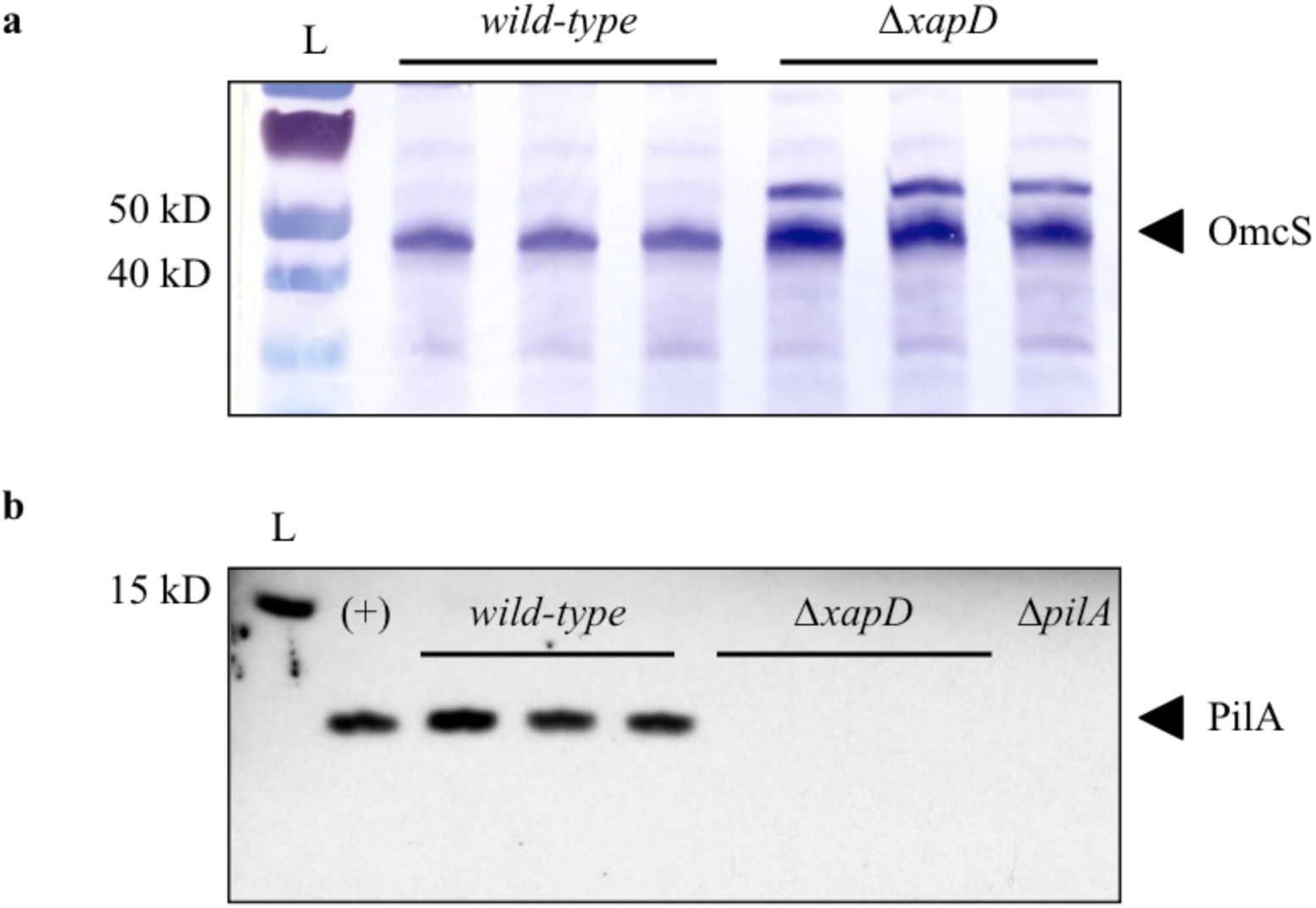
Western-blot analysis revealed presence of OmcS, but not PilA in the *xap-* deficient strain. A) Anti-OmcS Western blot analysis of loosely bound outer protein fraction, collected from triplicate mid-log cultures. B) Anti-PilA Western blot of whole cell lysates from triplicate, early stationary cultures of the wild-type and *xapD-deficient* strains of *G. sulfurreducens.* “(+)” and “Δ*pilA*” are positive and negative control samples, respectively; L = ladder.

However, we could not confirm that the *xapD*-deficient strain was producing PilA, the monomer necessary for pili production. No PilA was detected in whole cell lysates of the *xapD-*deficient strain with Western blots whereas identical conditions readily detected PilA in the wild-type strain (Fig. 1B).

### Localization of OmcS

It is difficult to understand how OmcS could be properly localized on e-pili if there is a lack of the PilA monomer to produce the pili. Therefore, in order to more definitively localize OmcS, the *xapD*-deficient and wild-type strains were examined with immunogold labeling with OmcS antibody (Fig. 2). Transmission electron microscopy of whole cell mounts of wild-type cells (Fig. 2A) revealed OmcS localized along pili, as previously reported^17^. In contrast, there was no immunogold labeling along pili in the *xapD-deficient* strain (Fig. 2B-D). Occasionally a few gold particles were observed, but they did not appear to be associated with pili (Fig. 2C), but rather near the outer surface of the cells (Fig. 2D).

**Figure 2.**
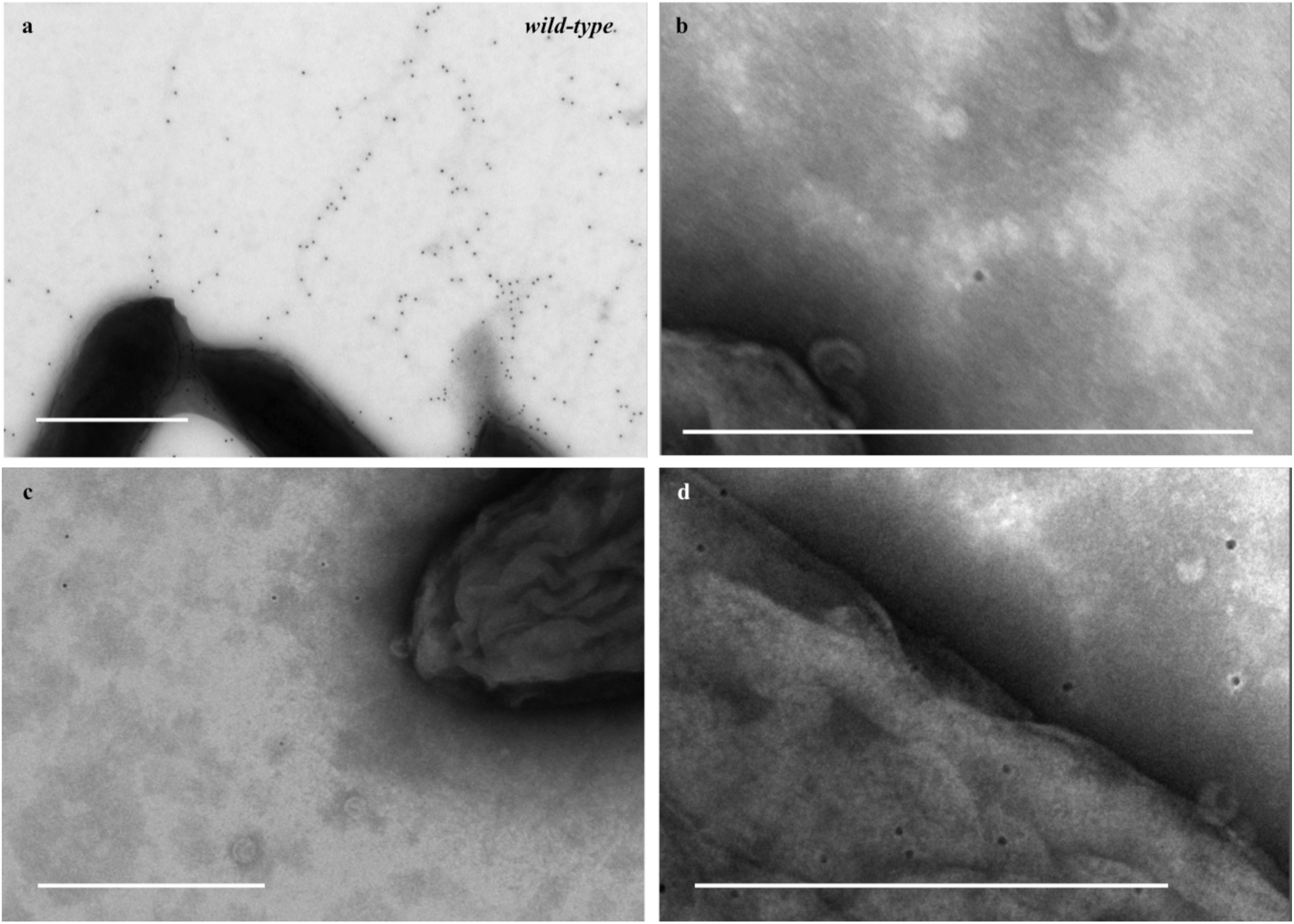
Transmission electron microscopy of immunogold-labeled whole cell preparations revealed that OmcS was associated with the pili in the wild-type strain of *Geobacter sulfurreducens,* but not in the *xapD*-deficient strain. The presence of OmcS is detectable as a uniform 10 nm electron-dense image. A) Wild-type strain with OmcS-decorated pili. B-D) Lack of OmcS-decorated pili in the *xapD-deficient* strain. Scale bars = 500nm.

In order to better define the localization of OmcS in the *xapD*-deficient strain, cells were further examined in ultrathin sections (Fig. 3). Pili are difficult to visualize with this method, but in wild-type cells the majority of the OmcS was detected at a distance from the cell, consistent with localization on pili (Fig. 2A). OmcS was also readily detected in *xapD*-deficient cells, but was closely associated with the cell surface (Fig. 3B). No OmcS was detected in an OmcS-deficient mutant (Fig. 3C), as previously reported^17^.

**Figure 3.**
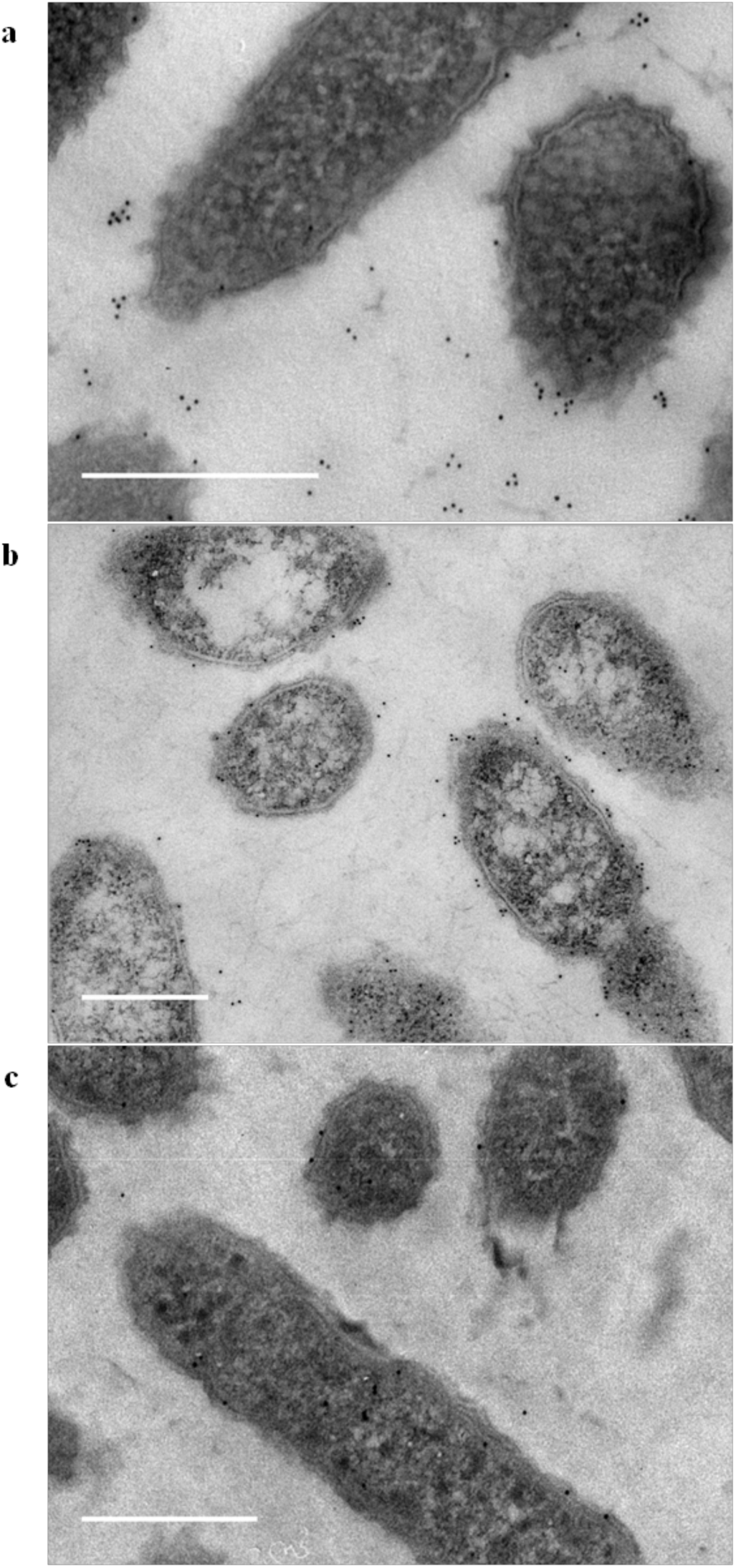
Transmission electron microscopy of immunogold-labeled thin sections revealed that OmcS was associated with outer cell surface of the *xapD*-deficient strain. A) Wild-type cells. B) *xapD-deficient* strain. C) *omcS*-deficient strain. Scale bars = 500nm.

## Implications

The results suggest that the previous conclusion^15^, based on indirect inference, that the *xapD*-deficient strain “still produced pili, as well as cytochromes attached to the pili”, is not correct. This is an important consideration in interpreting of the impact of deleting *xapD* on Fe(III) oxide reduction. Although it is clear that the *xapD* and related genes influence the expression of exopolysaccharide in *G. sulfurreducens*^15^, it also apparent that deleting *xapD* has the pleiotropic effect of mislocalizing OmcS and potentially inhibiting pili production. The finding that the *xapD*-deficient strain poorly reduces Fe(III) oxide even though there is abundant OmcS in the extracellular matrix emphasizes the importance of e-pili in properly positioning OmcS and is consistent with the concept^1,5^ that e-pili are a conduit for long-range extracellular electron transport and that OmcS facilitates electron transfer from e-pili to Fe(II) oxide.

Evaluation of the hypothesis that a cytochrome-rich outer-surface expolysaccharide matrix can also donate electrons to Fe(III) oxide, with an approach in which the expolysaccharide is diminished with gene mutations, will need to ensure that pleiotropic impacts on e-pili production and OmcS localization are avoided. Until pili formation in the *xapD-*deficient strain can be definitely demonstrated, claims for a role of the exopolysaccharide in attachment that is independent of pili participation also warrant scrutiny.

## Acknowledgements

We kindly thank Daniel Bond for the *xapD* deletion strain and Dale Callahan for TEM assistance.

## Author contributions

K.A.F., J.E.W. and D.R.L. designed experiments. K.A.F. ran Western blot analyses. K.A.F. and C.L. viewed whole-mount and ultrathin sections by TEM. J.E.W. conducted confirmatory experiments. K.A.F. and D.R.L. wrote the paper.

